# FLRT cell adhesion molecules counteract protocadherin-10-dependent cell segregation during cortical neuron migration

**DOI:** 10.64898/2026.07.26.740739

**Authors:** Yi-Ru Shen, Wenyu Ding, Daniel del Toro, SeungHee Chun, Tobias Straub, Gönül Seyit-Bremer, Fabian Coscia, Hao-Yi Li, Rüdiger Klein

## Abstract

The processes that underlie the folding of the cerebral cortex of large mammals and the evolution of the smooth cortex of modern rodents are incompletely understood. Recent evidence highlighted adhesion-controlled neuronal migration as an important mechanism. Genetic deletion in mice of FLRT1 and FLRT3 cell-adhesion proteins perturbed cortical neuron migration, lead to region-specific neuronal clustering, and introduced cortical folds into the normally smooth mouse cortex, suggesting that Increased expression of FLRT protein may have contributed to cortex smoothening. Here, we identify protocadherin-10 (PCDH10) as a key regulator of neuronal clustering in this context. Using single-cell transcriptomics and spatial proteomics, we describe a cortical neuron subtype in which loss of FLRT1/3 triggers local upregulation of PCDH10 in a specific cortical region. Elevated PCDH10 drives homophilic adhesion and the formation of cell patches that segregate from *FLR1/3*-deficient migrating neurons. This segregation leads to discrete clustering of *Flrt1/3*-mutant neurons and correlates with the emergence of cortical folds. Consistent with an evolutionary role, comparative expression analysis showed reciprocal expression of FLRT3 and PCDH10, with high FLRT3 and low PCDH10 expression in the smooth mouse cortex, and the opposite pattern in the developing human cortex. Together, our findings reveal how the antagonism between different classes of cell adhesion molecules affects cortical neuron migration patterns, contributes to region-specific cortical folding and may have promoted evolutionary smoothening of the rodent cortex.

## Introduction

The folding of the cerebral cortex into peaks (gyri) and fissures (sulci) has emerged in a highly stereotyped and evolutionarily conserved spatial pattern [1, 2]. In humans and other gyrencephalic species, major folds form at specific positions that often demarcate functional cortical areas, and are conserved across individuals and among phylogenetically related species [3–5]. Previous studies have shown that cortical thickness and folding are influenced by several factors, ranging from genetic and cellular programs to mechanical forces [6]. However, the mechanisms that specify folding patterns in specific spatial areas remain poorly understood.

Studies across gyrencephalic species indicate that primary cortical folds emerge in reproducible patterns predetermined by a transcriptomic protomap in cortical germinal zones. In ferret, thousands of genes involved in progenitor proliferation, neurogenesis, and cell fate specification are differentially expressed between the germinal zones of the prospective splenial gyrus and neighboring lateral sulcus. Often these genes distribute in modular blocks of high and low expression that anticipate the locations of future gyri and sulci [7]. Similar modular gene expression patterns are observed in macaque and human cortices but are absent in the lissencephalic mouse, supporting the existence of a genetic protomap underlying gyrencephalic folding patterns [7–10]. Consistent with this, subventricular zone (SVZ) modules that are thicker and more mitotically active preferentially give rise to expanding gyri, whereas thinner and less proliferative SVZ modules tend to generate sulci in ferret and macaque [11, 12]. This protomap is defined in part by differential enrichment of epigenetic marks such as H3K27ac, and block-wise expression of transcription factors involved in proliferation, adhesion, and migration [13].

Beyond proliferative patterning, cortical neuronal migration and lateral dispersion are another key mechanisms driving cortical folding [14], which act synergistically with progenitor expansion to shape gyrification [15]. Previous studies have shown that distinct neuronal subpopulations contribute differentially to cortical folding. For example, impaired migration of upper, but not lower-cortical neurons, following Cdk5 loss reduces cortical folding in the ferret [16]. These observations are supported by computational models, which suggest that distinct neuronal subpopulations differentially modulate biophysical parameters relevant to cortical folding, including tissue stiffness and cell density [17]. Moreover, lateral dispersion of neurons contributes to the expansion and folding of the cortex. Two major protein families, Eph/ephrins and FLRTs, have been shown to regulate lateral neuronal dispersion, and their knockdown or overexpression leads to neuronal clustering [18–20]. Interestingly, several studies have reported that such clustering precedes changes in cortical plate morphology, as neuronal segregation and heterogeneity give rise to features ranging from increased waviness [19] to cortical folding [14]. Other models of cortical folding have also described clustered neuronal distributions forming vertical column-like structures within the cortical plate [21].

We previously showed that *Flrt1/3* double knockout mice (dKO) develop sulci in latero-caudal regions of the cerebral cortex [14]. Cortex folding in these mutants depends on faster migration and cell clustering of *Flrt1/3*-mutant neurons in the cortical plate, which correlates with the formation of incipient sulcus in the mouse cortex. Notably, Flrt1/3 expression is much lower in the human neocortex than in the lissencephalic mouse, and reduced in prospective sulcal compared with gyral regions of the ferret cortex [13, 14]. Experimental data and modeling suggest that cell clustering in *Flrt1/3* dKO mice is non-cell-autonomous, raising the possibility that other molecular players participate in this process [14]. However, the identify of these mechanisms remains unknown.

Here, we combined single-cell RNA sequencing (scRNA-seq), and spatial proteomics to identify the molecular mechanism underlying neuronal clustering in *Flrt1/3* dKO mouse cortex. We found that about half of all cortical neurons expressed Flrt3 and that they were predominantly separated into two subpopulations distinguished by marker expression: an upper (Satb2+/Ctip2-) and a deeper (Satb2-/Ctip2+) population, which localize to distinct cortical regions. The latter is enriched in the latero-caudal cortex at E15.5, a region in which loss of FLRT1/3 induces neuronal clustering and sulcus formation specifically within this population. Mechanistically, the adhesion molecule protocadherin-10 (PCDH10) is selectively upregulated in FLRT3-negative neurons, which aggregate into patches through increased homophilic adhesion. This process drives the segregation and discrete clustering of *Flrt1/3*-mutant neurons, which correlates with sulcus formation. Together, these results support a role for adhesion-mediated cell sorting in regulating neuronal dispersion and in defining regional cortical morphology in specific spatial areas.

## Results

### scRNA-seq analysis reveals two distinct *Flrt3* neuron populations

We previously showed that loss of FLRT1/3 induces cell clustering in latero-caudal regions of the cerebral cortex at E15.5, which correlates with the formation of incipient sulci [14] (Figure 1A). To identify the molecular mechanisms underlying this region-specific phenotype, we first performed single-cell RNA sequencing (scRNA-seq) on latero-caudal cortex dissected from E15 *Flrt1/3* control (*Flrt1^−/−^*; *Flrt3^LacZ/+^*) and dKO (*Flrt1^−/−^*; *Flrt3^lox/LacZ^*; *Nestin-Cre*) embryos (Figure 1B). Transcriptomes from total 8,510 cells were plotted on Uniform Manifold Approximation and Projection (UMAP) (Figure 1C and Figure S1A-B). We assigned cell clusters into cell classes based on the coexpression of multiple marker genes, as done in previous studies [22] (Figure 1C and S1C-E).

**Figure 1.**
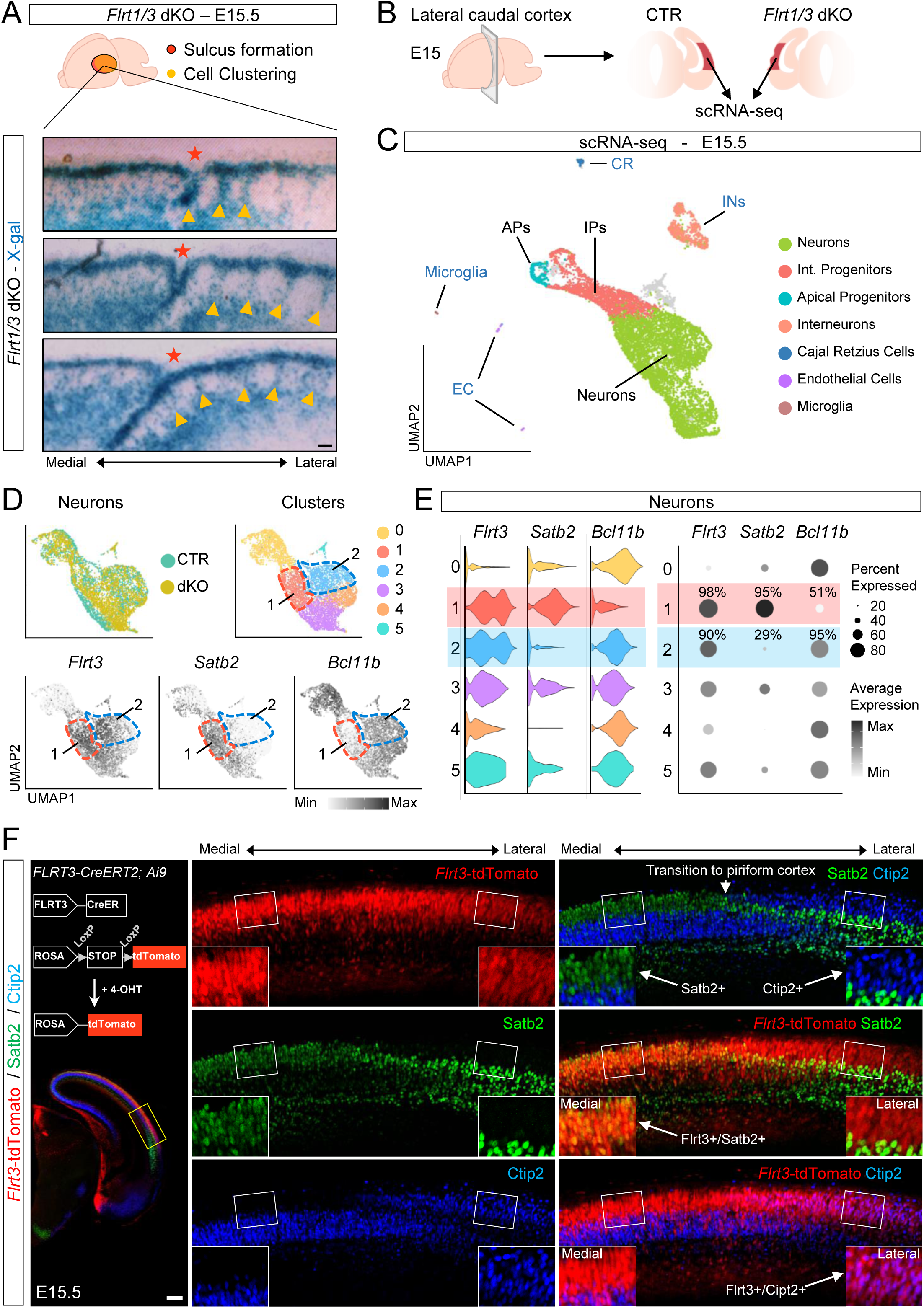
Two different FLRT3 neuron subtypes exhibit distinct spatial distribution patterns in the developing cortex. (A) Schematic (top) marks the locations of sulcus formation (red) and cell clustering (yellow) observed in E15 *Flrt1/3* dKO cortex. Serial coronal sections stained with X-gal from an E15 dKO brain reveal sulci (red stars) and clustered neurons (yellow arrowheads) in the lateral caudal cortex. Scale bar, 50 μm. (B) Schematic illustrating the dissected area of the caudal lateral cortex (dark red) from E15 control and dKO embryos for scRNA-seq. (C) UMAP with cells colored by cell types. Int. progenitors, intermediate progenitors. (D) UMAPs of excitatory neurons colored by genotype (left) and clustered subpopulations (right). Expression of *Flrt3*, *Satb2*, and *Bcl11b* in excitatory neurons is shown below. Clusters 1 and 2 are outlined in red and blue, respectively. (E) Violin plots and dot plots of *Flrt3*, *Satb2*, and *Bcl11b* expression across excitatory neuron clusters. Clusters 1 and 2 are outlined in red and blue, respectively. In dot plots, cell percentages for clusters 1 and 2 are shown above the corresponding dots. (F) Schematic of the inducible *Flrt3-CreERT2*; *Ai9* reporter system and staining of Satb2 (green), and Ctip2 (blue) in the caudal cortex of an E15 *Flrt3-CreERT2; Ai9* brain. Magnified medial-to-lateral views of the yellow box are at right. Scale bar, 300 μm.

Given that sulcus formation in *Flrt1/3* dKO cortices depends on clustering of migrating neurons [14], we focused our analysis on the neuronal population (5,025 cells), which segregated into 6 distinct clusters (cluster 0-5, Figure 1D). We analyzed the expression of *Flrt1* and *Flrt3* together with canonical neuronal markers across these clusters and found two distinct neuronal populations with high *Flrt3* expression (cluster 1 and 2) (Figure 1D-E and Figure S2A-B). Cluster 1 was characterized by high *Satb2* expression, with 95% of cells expressing *Satb2* and very low levels of *Bcl11b*/Ctip2. In contrast, cluster 2 showed an inverse expression pattern, with only 29% of cells showing low *Satb2* expression, while high *Bcl11b* expression was detected in 95% of cells (Figure 1E). We next asked whether *Flrt1/3* deletion altered the relative abundance of these subtypes. Although the exon 3 of *Flrt3*, containing the full coding sequence, was removed in dKO brains, reads mapping to exon 1 and 2 remained detectable, allowing the identification of *Flrt3* mutant cells (Figure S2C). Using this approach, we found that the proportions of the two subtypes did not change between control and *Flrt1/3* dKO cortices (Figure S2E-F). We next wanted to verify the presence and distribution of these two Flrt3-expressing neuronal populations *in vivo*. We crossed a Tamoxifen-inducible CreERT2 driven by the *Flrt3* promoter with a Cre-dependent tdTomato reporter (Ai9), generating the *Flrt3-CreERT2*;*Ai9* mouse line. This model allowed the visualization of FLRT3-expressing cells, as we previously showed [23]. We found that *Flrt3*-tdTomato cells were predominantly located in the cortical plate at E15.5, consistent with previous studies [14]. In medial regions, these cells were immunoreactive for Satb2, but negative for Ctip2. However, in lateral regions adjacent or encompassing the piriform cortex, these cells were negative for Satb2 and positive for Ctip2 (Figure 1F). Together, these results indicate the presence of two main subpopulations of FLRT3-expressing neurons: Flrt3+/Satb2+ and Flrt3+/ Ctip2+ that are enriched in distinct cortical areas along the medio-lateral axis.

### FLRT1/3 loss induces clustering exclusively in the Flrt3+/Ctip2+ neuron population

We previously showed that sulci in the *Flrt1/3* dKO line were predominantly located between the perirhinal and postrhinal cortices, close to the rhinal fissure [14]. The rhinal fissure is a highly conserved cortical sulcus that demarcates neocortex from piriform cortex in gyrencephalic species [24]. The observation that this region coincides with a boundary between two distinct FLRT3-expressing neurons (Satb2+ medial and Ctip2+ lateral), prompted us to quantify the distribution of these two neuron subtypes across the developing cortex.

We used the *Flrt3^LacZ/+^* reporter line to detect FLRT3-expressing cells by β-galactosidase (β-gal) staining, as previously described [14, 20]. Consistent with *the Flrt3-CreERT2;Ai9* reporter line, medial regions of caudal cortical sections were enriched in Flrt3⁺/Satb2⁺ neurons, whereas lateral regions were enriched in Flrt3⁺/Ctip2⁺ neurons (Figure 2A). Quantitative analysis across rostro-caudal levels showed that FLRT3-positive neurons were predominantly Satb2-positive in most cortical regions, whereas in the lateral caudal cortex, 84% of FLRT3-positive neurons lacked Satb2 expression (Figure 2C). Further analysis of this Satb2-negative FLRT3-positive population revealed that 81% of these cells expressed Ctip2 (Figure 2D). These findings are consistent with the scRNA-seq analysis identifying two distinct Flrt3 neuronal subtypes and further indicate that the Flrt3^+^/Satb2^-^/Ctip2^+^ subpopulation is selectively enriched in the lateral caudal cortex adjacent to the rhinal fissure.

**Figure 2.**
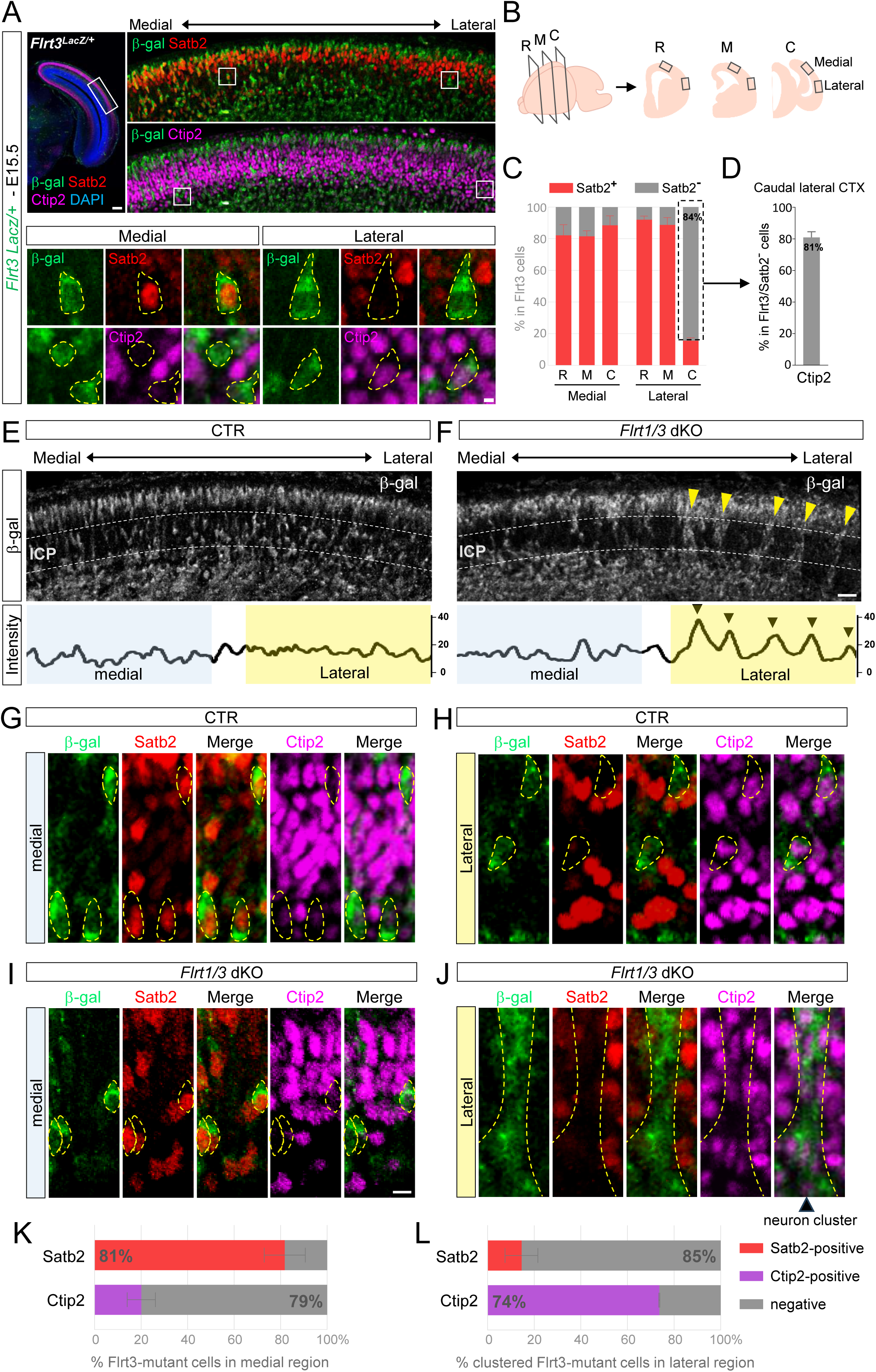
Loss of FLRT1/3 selectively drives clustering of Flrt3-deficient Satb2⁻/Ctip2⁺ neurons in lateral caudal cortex. (A) Immunostaining of β-galactosidase (β-gal, labels FLRT3^+^ neurons, green), Satb2 (red), and Ctip2 (magenta) in the caudal cortex of an E15 *Flrt3^LacZ/+^* embryo. Scale bar, 500 μm. Magnified images of the medial and lateral cortex are shown below. Yellow dashed outlines indicate β-gal+ cells. Scale bar, 24 μm. (B) Illustration of medial and lateral cortex in the rostral (R), middle (M), and caudal (C) levels. (C) Quantification of Satb2 expression among Flrt3^+^ neurons of E15 *Flrt3^LacZ/+^* brain in the CP of regions defined in (B). N=3 mice/group. Neurons>450/group. (D) Quantification of Ctip2 expression among Flrt3⁺/Satb2⁻ neurons in the caudal lateral cortex in (C). N=3 mice. Neurons>400. (E, F) β-gal staining (white) in the caudal cortex of E15.5 control (E) and *Flrt1/3* dKO (F) brains. Yellow arrowheads in (F) indicate clusters of FLRT1/3-deficient neurons. β-gal intensity plots for dashed rectangles are shown below. Black arrowheads mark intensity peaks corresponding to the FLRT1/3-deficient clusters. Scale bar, 50 μm. (G, H, I, J) Staining of β-gal (green), Satb2 (red), and Ctip2 (magenta) from medial and lateral regions of the control (G, H) and dKO (I, J) caudal cortex. Yellow dashed outlines highlight β-gal+ cells in G, H, I, and a neuron cluster in J. Scale bar, 12 μm. (K, L) Quantification of Satb2 and Ctip2 immunoreactivity among β-gal+ *Flrt3*-mutant neurons in *Flrt1/3* dKO cortices. Medial *Flrt3*-mutant neurons shown in (I) were quantified in (K), and lateral clustered *Flrt3*-mutant neurons shown in (J) were quantified in (L). Colored bars indicate Satb2-positive or Ctip2-positive cells, and gray bars indicate marker-negative cells. N = 3 mice; >350 neurons per group.

Next, we investigated whether these two distinct Flrt3 populations contribute to the discrete cell clustering observed in the *Flrt1/3* dKO cortex. We performed β-gal staining to identify *Flrt3*-mutant neurons (destined to express FLRT3) in *Flrt1/3* dKO embryos, and Flrt3+ neurons in control littermates in caudal cortical regions at E15.5. Consistent with previous studies [14], *Flrt3*-mutant neurons in dKO cortices were distributed in a repeated pattern of cell clusters within the lower cortical plate regions (Figure 2F). To quantify this distribution, we calculated intensity profiles of β-gal staining within the lower half of the cortical plate (dashed region), which revealed pronounced fluctuations in the density of *Flrt3*-mutant neurons compared with control FLRT3+ neurons in lateral but not medial regions (Fig. 2E–F). To pinpoint which subtype is involved, control cortices (Figure 2G–H) and dKO cortices (Figure 2I–J) were co-stained for β-gal, Satb2 and Ctip2. In *Flrt1/3* dKO cortices, the majority of clustered Flrt3-mutant (β-gal+) neurons in lateral regions were Satb2-negative (85%) and Ctip2-positive (74%) (Figure 2J and 2L). In contrast, *Flrt3*-mutant neurons that showed no evidence of clustering in medial regions were predominantly Satb2-positive and Ctip2-negative (Figure 2I and 2K). Therefore, loss of FLRT1/3 promotes neuronal clustering selectively within the Flrt3^+^/Satb2^-^/Ctip2^+^ subpopulation present in lateral regions.

### Loss of FLRT1/3 upregulates PCDH10 expression leading to homophilic adhesion

To identify the molecular mechanisms driving the discrete cell clustering of Flrt3^+^/Satb2^-^/Ctip2^+^ neurons in *Flrt1/3* dKO cortices, we performed spatial proteomic experiments using mass spectrometry coupled to laser microdissection. We used X-gal staining to visualize mutant neurons in *Flrt1/3* dKO cortices, allowing microdissection of cortical plate regions showing cell clustering, as well as adjacent control areas, such as the intermediate zone, where mutant neurons did not cluster (Figure 3A). We then compared the proteomic profiles from these regions with corresponding areas from control (CTR) cortices.

**Figure 3.**
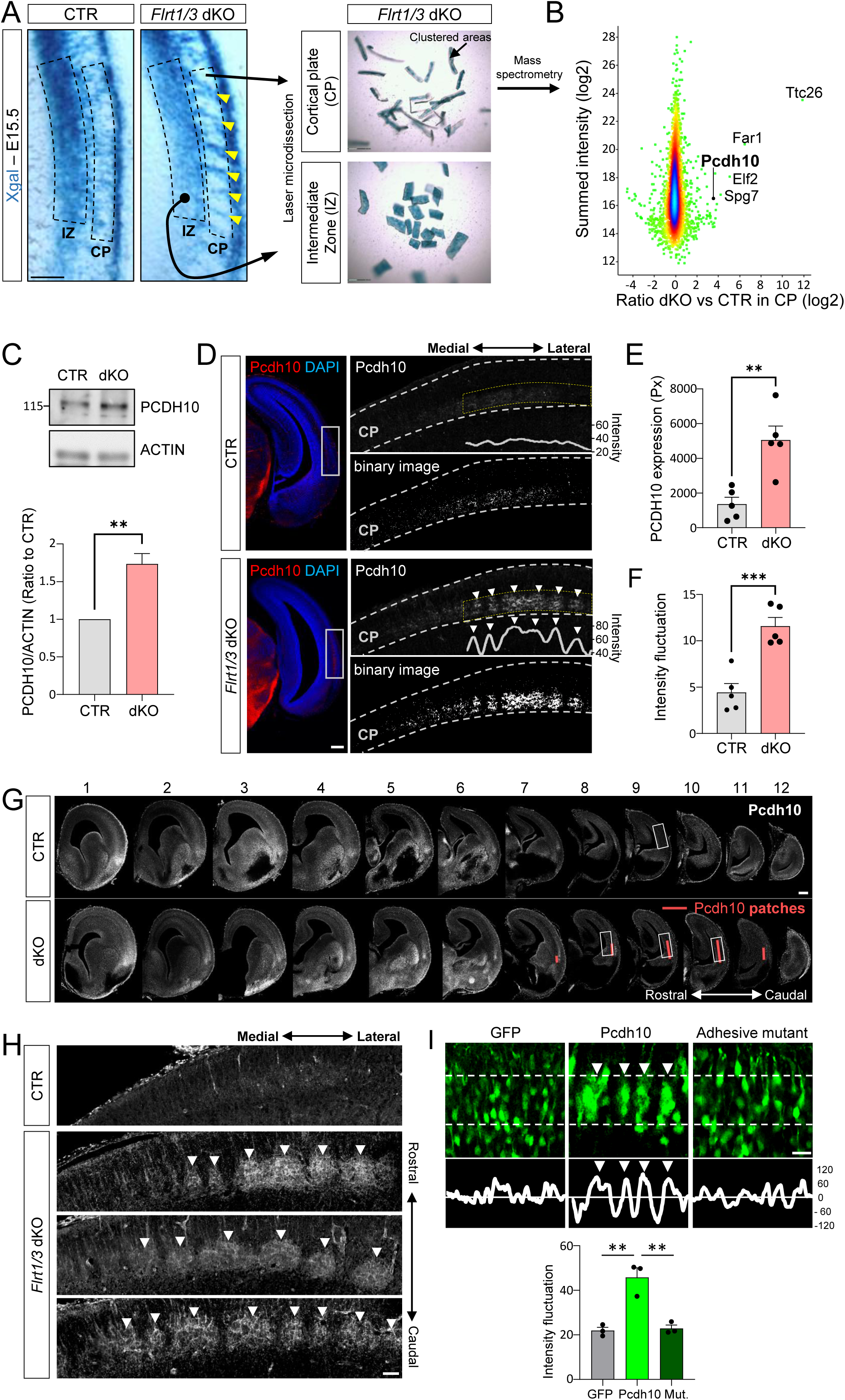
Spatial proteomic reveals increased PCDH10 expression and cell patch formation in *Flrt1/3* dKO cortex. (A) X-gal staining of coronal sections from E15 control and *Flrt1/3* dKO caudal brains highlights Flrt3 neurons in blue. Yellow arrowheads indicate clusters of *FLRT1/3*-deficient neurons. Dashed lines outline the CP and IZ regions of lateral-caudal cortices collected by laser microdissection. Scale bar, 150 μm. Dissected CP (clustered) and IZ (non-clustered) areas of dKO cortices are shown at right. (B, C) Volcano plots for CP show normalized protein expression ratios (dKO vs. CTR, log₂) plotted against summed intensity (log₂). The points are colored by their “significance B”, with blue points having values of 0.5 or greater, red between 0.1 and 0.5, yellow between 0.05 and 0.1 and green below 0.05. (C) Western blot of E15 control and dKO caudal cortex using antibodies against PCDH10 and ACTIN. Quantification of PCDH10/ACTIN signal ratio from three biological replicates is shown below. (D) Immunofluorescence of PCDH10 in E15 control and dKO brain sections. Higher magnification of white rectangle is shown on the right. Arrowheads indicate PCDH10-positive cell patches. Scale bar, 500 μm. (E) PCDH10-positive area of CP regions in (D) were quantified from binary images thresholded in ImageJ (pixel intensity >70). Px, square pixels. N=5 mice/group. (F) Quantification of PCDH10 intensity fluctuations across yellow boxes (intensity plots are shown below) in (D). N=5 mice/group. (G, H) Serial coronal sections (1-12) from an E15 *Flrt3^LacZ/+^* brain, arranged from rostral to caudal, showing PCDH10 expression (white). Red bars mark PCDH10 patches (G). Scale bar, 1 mm. Magnified views of boxed areas are shown in (H). Arrowheads indicate PCDH10 patches. Scale bar, 50 μm. (I) Wild-type cortices electroporated with GFP, Pcdh10-GFP, or adhesion-deficient mutant. Arrowheads indicate PCDH10 patches. Scale bar, 25 μm. Intensity profiles across the white rectangle and quantification of intensity fluctuations are shown below. N=3 mice/group. **p < 0.01, ***p < 0.001, two-tailed Student’s t-test.

This analysis identified several proteins that were increased in the dKO cortical plate, representing diverse cellular processes, including ciliary transport (TTC26), lipid metabolism (FAR1), transcriptional regulation (ELF2), mitochondrial function (SPG7), and cell-surface adhesion (PCDH10) (Figure 3B and Table S1). We focused on PCDH10 because its established homophilic adhesive activity made it the most compelling candidate to regulate neuronal segregation in the *Flrt1/3*-mutant cortex [25]. In addition, the upregulation of PCDH10 was restricted to the clustering regions of the cortical plate and was not observed in the intermediate zone (Figure S3A). We confirmed this upregulation by both western blot (Figure 3C) and quantitative immunostaining (Figure 3D-E). These findings were consistent with our scRNA-seq analysis of neurons from control and dKO cortices, which identified 503 differentially expressed genes (DEGs), including several receptors involved in cell adhesion such as PCDH10 (Figure S3B and Table S1).

We next analyzed the distribution of cells expressing PCDH10 in caudal cortical sections by examining their intensity profiles. In contrast to control sections, which showed a weak and uniform PCDH10 staining, *Flrt1/3* dKO sections showed a strong and patchy distribution with high fluctuations (Figure 3D-F). Analysis of 12 serial brain sections along the rostral-caudal axis showed that PCDH10+ cells distributed in patches that were confined to the lateral caudal region of *Flrt1/3* dKO cortices at E15 (Figure 3G-H). Given that PCDH10 is a homophilic cell-cell adhesion molecule [25], we hypothesized that elevated PCDH10 expression could drive cell aggregation and the formation of patchy domains observed in *Flrt1/3* dKO cortices. We generated pCAG vectors encoding PCDH10 and an adhesion-deficient PCDH10 mutant by introducing a single point mutation (L105K) in the extracellular cadherin repeat 1, as described previously [26]. We first used these constructs in cell aggregation assays with K-562 cells. Consistent with previous studies, ectopic expression of wild type PCDH10 induced cell aggregation (Figure S3C-D), whereas expression of the PCDH10 L105K mutant did not (Figure S3E). We next performed in vivo gain-of-function experiments using *in utero* electroporation, following our established approach for FLRT proteins [20]. Overexpression of wild-type PCDH10, but not the adhesion-deficient L105K mutant, induced neuronal aggregation and formation of patchy domains within the cortical plate (Figure 3I). These results indicate that high PCDH10 expression is sufficient to induce neuron aggregation through homophilic trans-interactions in the cortical plate during cortex development.

### *Flrt1/3* mutant neurons segregate from PCDH10+ patches

Given that PCDH10+ cells displayed a patchy distribution whereas *Flrt1/3* mutant cells formed discrete clusters, we next examined the spatial relationship between these two populations. We performed immunostaining to visualize PCDH10+ cells in *Flrt1/3* dKO cortices together with β-gal staining of mutant cells carrying *Flrt3*-LacZ reporter allele. Intensity fluctuation analysis showed that *Flrt1/3* mutant neurons segregated from PCDH10+ patches and formed discrete clusters (Figure 4A). This pattern was not observed in control cortices or in medial regions of *Flrt1/3* mutant cortices, where PCDH10 protein levels were low and FLRT1/3+ cells were uniformly distributed (Figure S4A-E). Moreover, the *Flrt1/3* mutant neurons that segregated from PCDH10+ patches corresponded to the *Flrt3*-knockout Satb2-negative neuronal subpopulation (Figure S4B and S4F).

**Figure 4.**
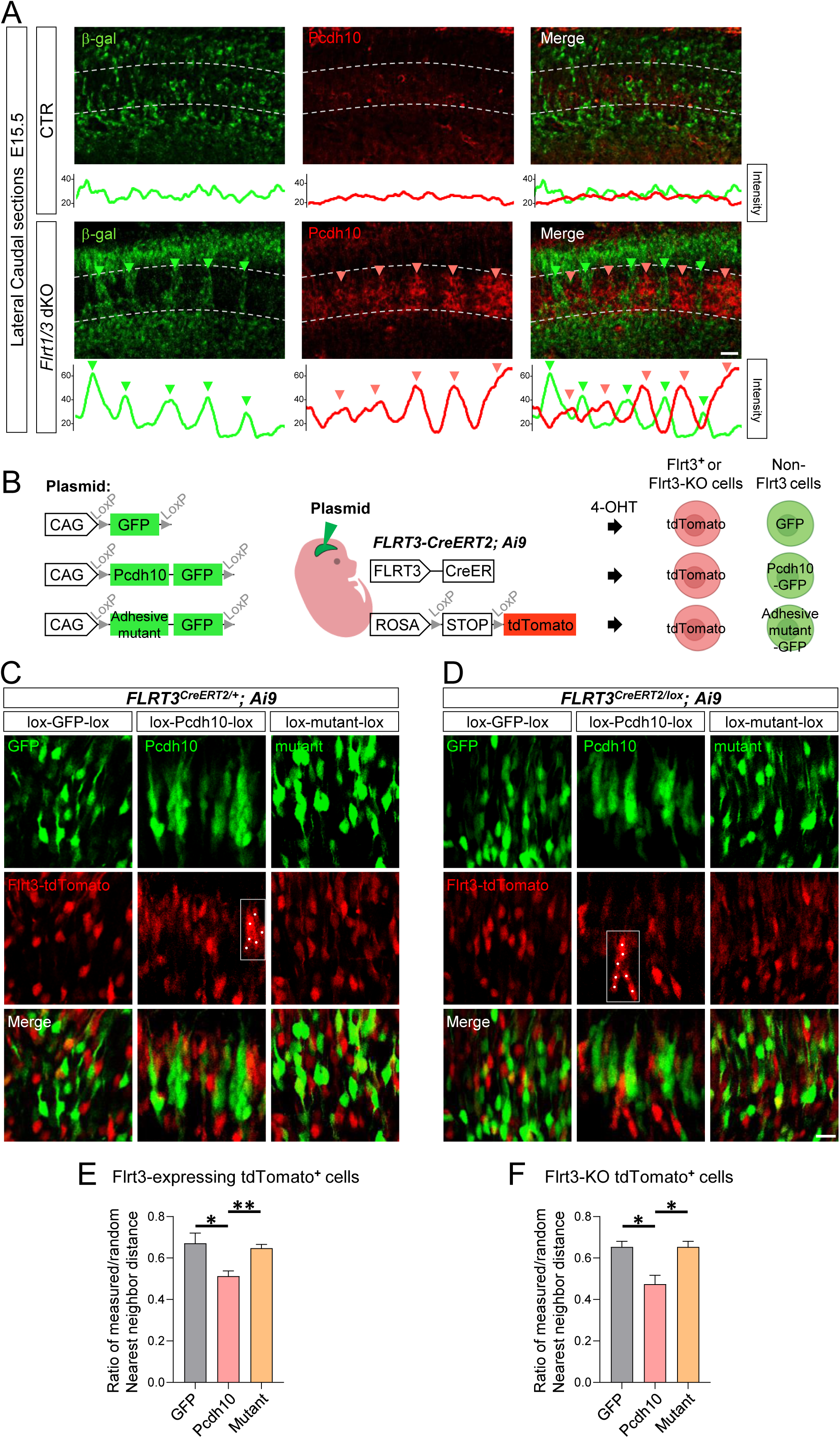
PCDH10-mediated adhesion contributes to segregation between PCDH10 cell patches and Flrt3 neurons. (A) Immunostaining of β-gal (green) and PCDH10 (red) in the caudal lateral cortex of E15.5 control and *Flrt1/3* dKO brains. Intensity profiles across the white rectangles are shown below. Green and pink arrowheads mark FLRT1/3-deficient clusters or PCDH10 patches and their corresponding intensity peaks, respectively. Scale bar, 50 μm. (B) Schematic of plasmids used for *in utero* electroporation (IUE) and the *Flrt3-CreERT2*;*Ai9* strategy. Plasmids flanked by loxP sites were electroporated into cortices of *Flrt3*-*CreERT2*; *Ai9* embryos, in which CreERT2 expression is driven by the *Flrt3* promoter and tdTomato expression is Cre-dependent. Following 4-OHT induction, CreERT2 activity in Flrt3 cells excised the loxP-flanked cassette and simultaneously activated tdTomato expression. Consequently, Flrt3 neurons expressed tdTomato, whereas non-FLRT3 expressing wild-type (non-Flrt3) cells expressed the loxP-flanked cassette. (C, D) IUE of GFP, Pcdh10-GFP, or adhesion-deficient mutants in *FLRT3^CreERT2/+^; Ai9* (C) and *FLRT3^CreERT2/lox^; Ai9* (D) cortices. White boxes highlight clusters of Flrt3-tdTomato neurons; white dots mark individual cells. Scale bar, 25 μm. (E, F) Ratio of measured to theoretical random nearest-neighbor distances between Flrt3-expressing tdTomato⁺ cells (E) and Flrt3-deficient tdTomato⁺ cells (F). Measured values represent the actual distance to the nearest tdTomato⁺ neighbor; random distances are calculated assuming uniform spatial distribution. A lower ratio indicates stronger clustering. N = 3 mice per group. *p < 0.05, **p < 0.01, two-tailed Student’s t-test.

These results raised the possibility that the increased PCDH10 expression and cell aggregation in FLRT-negative neurons in the *Flrt1/3* dKO cortex were sufficient to promote segregation and clustering of *Flrt1/3*-mutant neurons in a process of differential cell adhesion. To test this, we designed a strategy using the *Flrt3-CreER;Ai9* reporter mouse line to differentially target FLRT3-expressing and FLRT3-negative neurons (Figure 4B). Using Cre-dependent vectors, we induced PCDH10 expression together with GFP selectively in neurons that do not express FLRT3 (CreER -negative), which represented ∼50% of migrating neurons at E15.5, as shown previously [14, 15]. Instead, in *Flrt3*-CreER-expressing neurons the floxed PCDH10-GFP expression cassette was excised and cells were labeled by Cre-dependent tdTomato expression from the Ai9 reporter line. We validated this strategy by electroporating GFP control plasmids with or without loxP flanking sites into *Flrt3-CreER;Ai9* embryos (Figures S4G-H). The non-floxed GFP plasmid was expressed in tdTomato-positive Flrt3 neurons, resulting in numerous double-positive cells. In contrast, Cre-mediated excision of the floxed GFP cassette in FLRT3-expressing neurons restricted GFP expression to the non-FLRT3 population, reducing double-positive cells to ∼10%. These results confirmed that this strategy differentially targets FLRT3-expressing and FLRT3-negative neurons.

Next, we examined the effects of expressing PCDH10 or its adhesion-deficient L105K mutant specifically in FLRT3-negative cells *in vivo*. We did this both in heterozygous *Flrt3-CreERT2/+*; *Ai9* (Figure 4C) and FLRT3-null *Flrt3CreERT2/lox*; *Ai9* (Figure 4D) embryos to determine whether the observed effects depended on FLRT3 protein expression. FLRT3-negative cells expressing wild-type, but not mutant, PCDH10 formed prominent aggregates that segregated from FLRT3-expressing tdTomato neurons. This happened in both control (Figure 4C) and FLRT3-deficient (Figure 4D) cortices, indicating that segregation between PCDH10 cell aggregates and Flrt3 neurons was driven by PCDH10 expression and occurred independently of FLRT3 expression. Notably, the distance between neighboring *Flrt3*-tdTomato neurons was significantly reduced following PCDH10 overexpression compared with GFP controls (Figure 4E-F), indicating that elevated PCDH10 expression reduced the distance between Flrt3 neurons, promoting their clustering. In contrast, neurons expressing the adhesion-deficient PCDH10 mutant were evenly distributed and did not affect the distance between tdTomato-positive cells (Figure 4C-F). Together, these results suggest a model in which PCDH10-mediated cell adhesion of FLRT3-negative neurons drives cell segregation and subsequent clustering of Flrt3+ neurons by differential cell adhesion.

### Modeling clustering of *Flrt1/3* dKO neurons by PCDH10-induced segregation

To further validate this model, we characterized the neuronal population expressing PCDH10 by co-staining with cortical neuron markers. We found that PCDH10 was upregulated and formed patches specifically in the lower part of the cortical plate in lateral caudal regions of *Flrt1/3* dKO brains at E15, but not earlier developmental stages (Figure 5A-B). This temporal and spatial pattern correlated with the emergence of *Flrt1/3* mutant cell clusters, as shown previously [14] (Figure 5B). Moreover, PCDH10+ cells within these patches expressed deeper layer cortical markers (Tbr1+) at the time when *Flrt1/3* mutant neurons were migrating through this layer to reach the upper part of the cortical plate [14](Figure 5B). This result is consistent with our scRNA-seq data where ∼80 % of Pcdh10-expressing cells were Tbr1⁺ neurons (Figure S5A).

**Figure 5.**
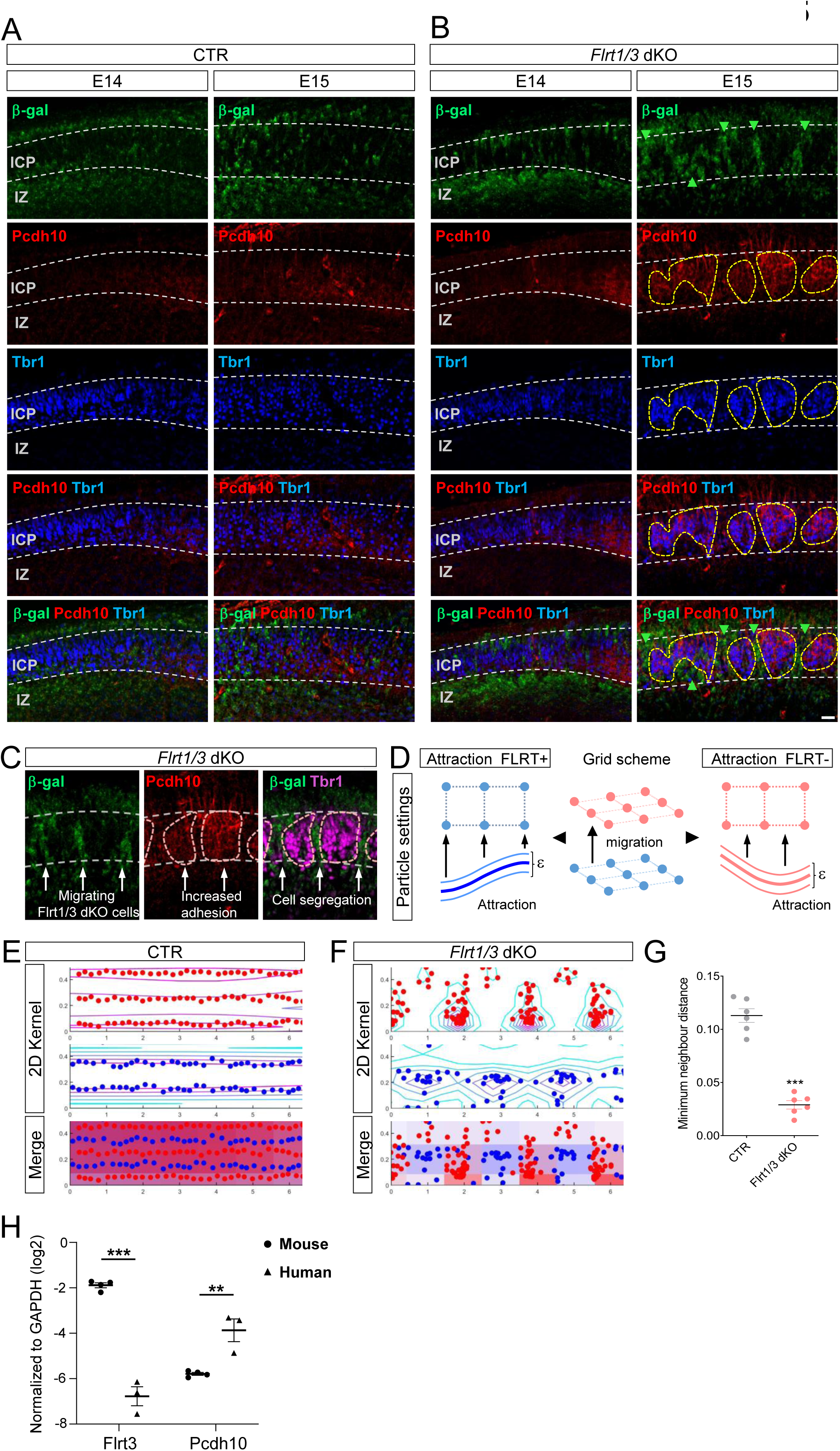
Simulation suggests that PCDH10-mediated differential adhesion sorts migrating FLRT1/3-deficient cells to cluster in the spaces between PCDH10 patches. (A, B) Immunostaining of β-gal (green), PCDH10 (red), and Tbr1 (magenta) in the lateral caudal cortex of E14 and E15 control (A) and *Flrt1/3* dKO (B) embryos. Green arrowheads indicate clusters of FLRT1/3-deficient neurons. Yellow dashed outlines highlight PCDH10 and Tbr1 patches. Scale bar, 50 μm. (C) Migrating *Flrt1/3* mutant neurons (β-gal⁺, green) segregate from PCDH10⁺/Tbr1⁺ neurons (red/magenta) located in the lower cortical plate (dashed lines). (D) Scheme illustrating how particles representing FLRT1/3-positive (blue) and -negative neurons (red) are arranged. Blue and red particles show attraction between themselves, modeled as a sinus equation. ε represents noise added to the system. (E-F) Distribution of particles representing FLRT1/3-expressing (blue) and non-expressing (red) neurons after computer simulations under control (CTR) and *Flrt1/3* dKO conditions. Single channels show in colored lines higher (magenta) or lower (cyan) density of particles based on their kernel distribution. Merge images show both particles together with their density as blue and red filled areas. (G) Minimum neighbor distance of particles shown in (E-F). n = 10 computer simulations comprising 204 (red) and 165 (blue) particles per run. ∗∗∗p < 0.001, one-way ANOVA with Tukey’s post hoc test. (H) Comparison of FLRT3 and PCDH10 expression levels between mouse and human with bulk RNaseq data. E15 mouse embryonic cortices from Nitarska, et al. (GEO: GSE70298) [28] were compared to early mid-trimester fetuses at 16 pcw from the Allan Brain Atlas (http://www.brain-map.org). Values were log-transformed and expressed relative to GAPDH. **p < 0.01, ***p < 0.001, two-tailed Student’s t-test.

These results support a two-layer model in which Tbr1+ neurons in the lower cortical plate upregulate PCDH10 leading to their aggregation into patches, while migrating *Flrt1/3* mutant neurons entering from the intermediate zone are segregated and cluster between these patches (Figure 5C). We adapted our previous data-driven computational modeling of FLRT1/3+ neurons migrating through the cortical plate [14]. In this model, we considered an upper layer of neurons through which FLRT1/3+ neurons must pass, which showed either low adhesion between cells (control condition) or high adhesion due to PCDH10 upregulation in the *Flrt1/3* dKO condition. Adhesive forces were modeled as sine equations following the repeated pattern of cells in the *Flrt1/3* dKO condition (Figure 5C), as done previously [14](Figure 5D). To analyze the behavior of the moving particles, FLRT-positive particles were set to move along the z axis, and both speed and attraction forces were random within a small range (ε) to mimic fluctuations present in biological systems [27]. In control conditions, with low attraction among FLRT1/3-positive and negative particles, the particles showed a homogeneous distribution after moving along the z axis (Figure 5E). In the *Flrt1/3* dKO situation, FLRT1/3-negative particles aggregated into patches due to high attraction (mediated by PCDH10), whereas *Flrt1/3* mutant particles segregated and formed clusters between these patches (Figure 5F-G). Together, these simulations recapitulate the experimental observations, showing that increased adhesion among FLRT1/3-negative cells is sufficient to segregate and cluster *Flrt1/3*-mutant neurons.

Finally, we asked whether the reciprocal high FLRT3 and low PCDH10 expression pattern observed in the normal mouse cortex was inverted in the developing human cortex as it was in the *Flrt1/3* dKO mouse model. We compared FLRT3 and PCDH10 mRNA expression levels in E15 mouse neocortex [28] with RNA-seq data from human fetal cortex at 16 post-conception weeks (pcw) (http://www.brain-map.org), normalized to the housekeeping gene GAPDH (Figure 5H). FLRT3 expression was higher in embryonic mouse cortex than in human fetal cortex, consistent with our previous findings [14]. Interestingly, PCDH10 showed the opposite pattern, with higher expression in human fetal than embryonic mouse cortex (Figure 5H). These results revealed that the expression patterns of FLRT3 and PCDH10 are inverted in the human cortex and more closely resemble the *Flrt1/3* dKO than the normal mouse cortex.

## Discussion

In this study, we identified a cell-adhesion based mechanism that promotes region-specific neuronal clustering associated with cortical folding. We previously showed that genetic deletion of FLRT1/3 receptors alters neuronal migration, leading to discrete cell clustering that correlates with sulcus formation without progenitor amplification [14]. Here, we showed that this phenotype was driven by increased PCDH10-mediated homophilic adhesion, which promoted spatial segregation of two distinct neuronal subpopulations that correlated with sulcus formation in laterocaudal cortical regions (Figure S5B). Mechanistically, this clustering was characterized by selective upregulation of PCDH10 in Tbr1+ neurons within the lower cortical plate, which assembled into patches through PCDH10-homophilic adhesion. These PCDH10+ domains segregated migrating *Flrt1/3* mutant neurons, mainly Satb2^-^/Ctip2^+^neurons, resulting in their discrete clustering as they migrated through the cortical plate. Thus, the study reveals an antagonistic balance between FLRT and PCDH10-mediated cell adhesion that regulates neuronal spatial organization and cortical folding.

### PCDH10 as a mechanism underlying cell clustering in *Flrt1/3* dKO cortices

In *Flrt1/3* dKO embryos, we found that PCDH10 was upregulated in Tbr1+ neurons present in laterocaudal cortical regions. Our experimental and modeling analysis suggest that this upregulation is sufficient to drive aggregation of these cells into patches, which in turn segregate migrating *Flrt1/3*-mutant neurons into discrete clusters. This mechanism is consistent with the established role of PCDH10 in trans-homophilic adhesion during circuit assembly and synaptic maintenance [26, 29–33], as well as with the differential adhesion hypothesis. In this framework, highly adhesive PCDH10+ cells form cohesive patches, whereas less adhesive Ctip2+ and *Flrt1/3*-mutant neurons are segregated into clusters [34]. Such differential adhesion–mediated sorting would generate an alternating pattern, consistent with both our *in vivo* observations and computational modeling. Moreover, this mechanism provides a satisfactory explanation for our previous findings, in which experimental data and modeling had suggested that FLRT1/3-deficient neurons were repelled by surrounding cells [14]. The mechanisms underlying PCDH10 upregulation in the absence of FLRT1/3 are not known. FLRT receptors are known to form multimolecular complexes with partners such as Latrophilins, Teneurins, and Unc5 receptors, which regulate neuronal activity, adhesion, and cytoskeletal dynamics [20, 35, 36]. Alterations in these processes could indirectly influence PCDH10 expression, as its expression has been shown to respond to activity-related signaling in developing olfactory sensory neurons [37]. In addition, changes in tissue mechanics and cellular tension are known to regulate adhesion gene programs, including cadherins, suggesting that altered mechanical properties of the cortex in *Flrt1/3* mutants could further contribute to PCDH10 upregulation [38]. How FLRTs could influence the expression of PCDH10 or other cell adhesion receptors is a stimulating question for future research.

Protocadherin-driven cell sorting has been observed in diverse morphogenetic processes, including cortical folding. In the developing cortex, mosaic PCDH19 expression caused by random X-inactivation drives differential adhesion that segregates wild-type and mutant clones, leading to aberrant gyrification [39, 40]. Differential PCDH19 expression also delineates lineage boundaries in the zebrafish spinal cord and maintains the entorhinal–neocortical interface in mice [41, 42]. Similarly, during Xenopus gastrulation, complementary expression of axial Pcdh1 and paraxial Pcdh8 confers distinct adhesion properties that partitions the mesoderm into axial and paraxial domains [43, 44]. Together with our PCDH10 data, these findings highlight protocadherin-mediated differential adhesion in tissue morphogenesis, including cortical folding.

### Evolutionary considerations

Previous genetic mouse models in which cortical folding is driven by progenitor amplification typically show folds at stochastic or random locations across the cerebral cortex [8, 9, 45, 46]. In contrast, the *Flrt1/3* dKO mouse model shows a spatial preference, with sulcus formation occurring preferentially between the perirhinal and postrhinal cortices [14]. This region, close to the rhinal fissure, corresponds to a cortical area in which sulcus formation is highly conserved in gyrencephalic species [24]. This spatial specificity is not mirrored by the overall expression patterns of Flrt1, which is largely uniform across the cortex, or by Flrt3, which shows a gradual rostro-caudal and medio-lateral gradient [14, 35]. Instead, it correlates with the distribution of a specific neuronal subpopulation that coexpresses Flrt3 and Ctip2 in laterocaudal regions of the cortex, suggesting that it may be particularly sensitive to the loss of FLRT1/3. This interpretation is consistent with previous studies showing differential sensitivity of piriform and neocortical neurons to genetic perturbations such as loss of Lhx2 and Mecp2 [47, 48]. In line with this, glutamatergic neurons in the piriform cortex show distinct developmental and transcriptomic programs compared to neocortical neurons [49], potentially reflecting differences in their projection targets [50]. Moreover, Ctip2+ neurons show higher expression and distinct spatial organization in the piriform cortex compared to the neocortex [51, 52]. In future work, it would be interesting to assess whether there are specific FLRT-dependent protein complexes in the piriform cortex compared with other cortical areas. Recruitment of distinct binding partners may activate specific intracellular signaling pathways in neighboring cells, potentially regulating the expression levels of other receptors in addition to PCDH10. Indeed, the finding that the gyrencephalic human cortex expresses higher levels of PCDH10 than the lissencephalic mouse cortex, whereas Flrt1/3 expression shows the opposite pattern and is much lower in human cortex relative to mouse (Figure 5H)[14], suggests a potential regulatory mechanism between these two systems. In this framework, increased FLRT expression in the mouse prevents PCDH10-dependent neuronal segregation contributing to a smooth cortex, whereas lower FLRT expression in gyrencephalic species favours this process, increasing neuronal lateral dispersion and promoting stereotyped cortical folding.

Our findings thus reveal an antagonism between two classes of cell adhesion molecules, FLRT and PCDH10, that regulates cortical neuron migration patterns and contributes to region-specific cortical folding. We identify PCDH10 as a key mediator of neuronal segregation that becomes selectively upregulated in Tbr1+ neurons of the laterocaudal cortex following loss of FLRT1/3, leading to the formation of adhesive patches that segregate migrating FLRT1/3-deficient neurons. A similar expression pattern is observed in the developing human neocortex, which expresses much lower levels of Flrt1/3 and higher levels of PCDH10 than the mouse cortex. This scenario provides a molecular and cellular framework linking differential cell adhesion to neuronal lateral dispersion, and suggests that changes in the balance between FLRT-and PCDH10-mediated adhesion may have contributed to the evolution of neuronal migration from gyrencephalic to lissencephalic species.

## Acknowledgements

We thank the image facility of the Max Planck Institute for Biological Intelligence (Martinsried, Germany), and the proteomics facility and the next generation sequencing facility of the Max Planck Institute of Biochemistry (Martinsried, Germany).

## Author contributions

Y.-R.S., D.dT and R.K. conceptualized the study and designed experiments. Y.-R.S. performed most of the experiments. W.Y.D. contributed to scRNA-seq. S.H.C. performed surgeries. T.S. helped with scRNA-seq data. G.S.B contributed with mouse experiments. F.C. processed LC-MS/MS analysis. H.-Y.L. performed bulk RNaseq analysis. D.d.T. performed computational simulations; R.K. and D.d.T. supervised; Y.-R.S., D.d.T. and R.K., wrote the manuscript with help from all authors; R.K. provided funding.

## Figure legends

**Supplementary figure 1.**
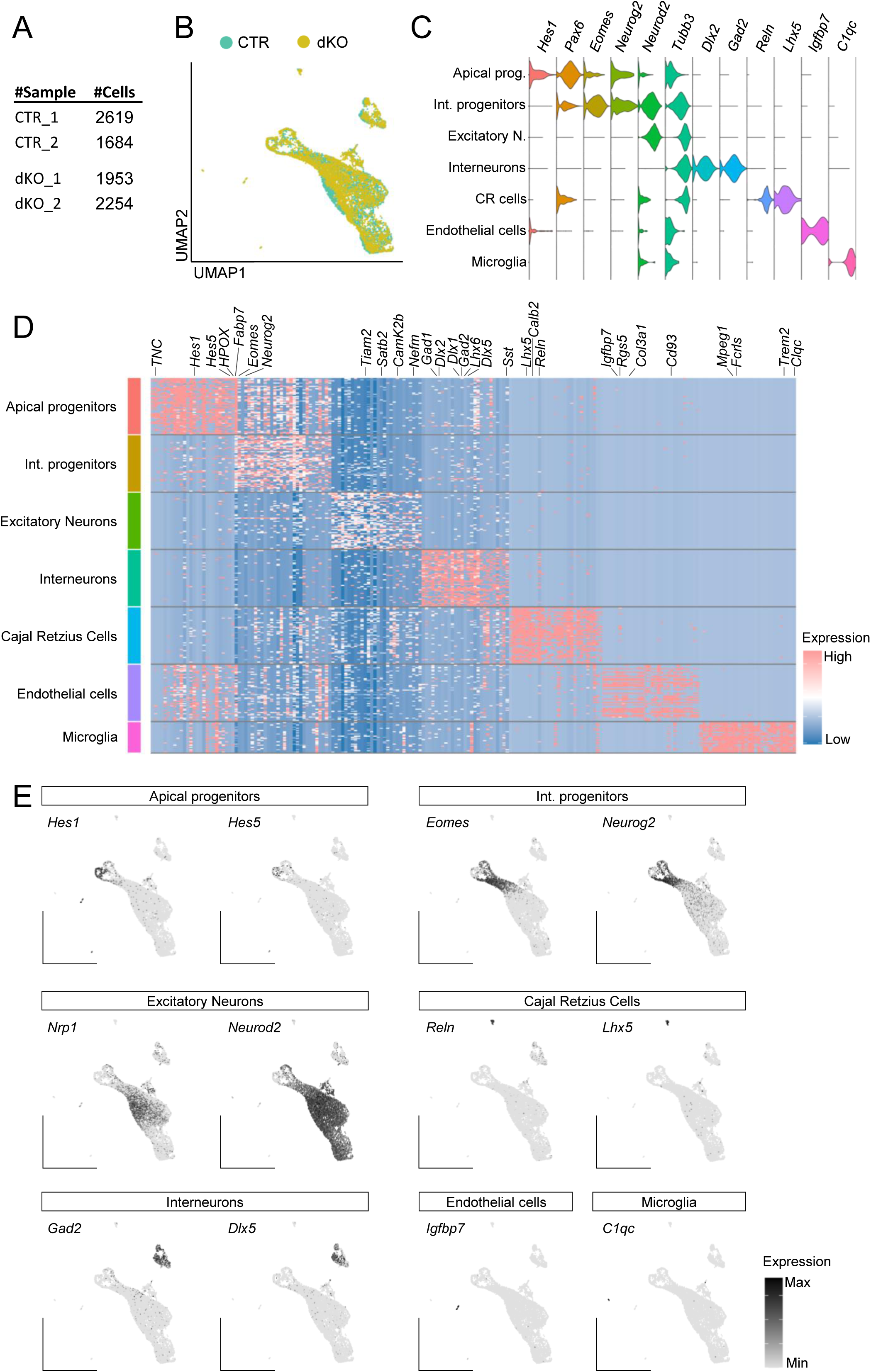
Molecular signatures defining cell types in scRNA-seq analysis. (A) Table summarizing the number of cells obtained per replicate from control and *Flrt1/3* dKO samples. (B) UMAP with cells colored by genotype. (C) Violin plots with expression of canonical marker genes for each cell type. (D) Heatmap of the top 30 differentially expressed genes for each cell type. Cells were downsampled to a maximum of 50 per cell type. (E) UMAP with expression of canonical marker genes for each cell type.

**Supplementary figure 2.**
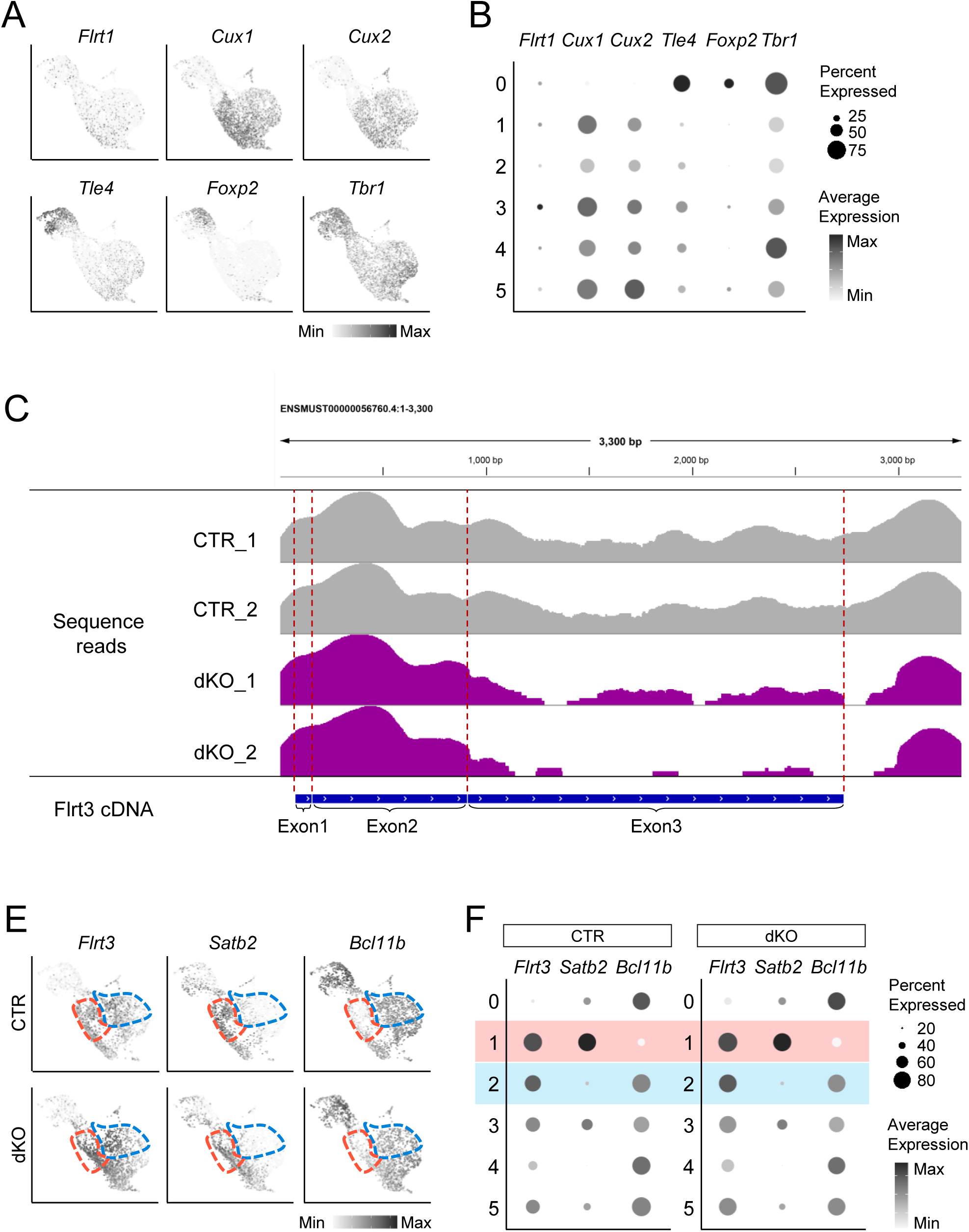
Molecular features of excitatory neuron subtypes in control and *Flrt1/3* dKO cortex. (A, B) UMAP (A) and dot plots (B) showing expression of *Flrt1* and layer-specific markers (*Cux1*, *Cux2*, *Tle4*, *Foxp2*, *Tbr1*). (C) Alignment of scRNA-seq reads mapped to the Flrt3 cDNA. In dKO samples, reads corresponding to *Flrt3* exons 1 and 2 remain detectable, whereas coverage over exon 3 is greatly reduced, consistent with Cre-mediated deletion of the coding region. (E, F) UMAP (E) and dot plots (F) of *Flrt3*, *Satb2*, and *Bcl11b* expression in control and dKO samples. Clusters 1 and 2 are outlined in red and blue, respectively.

**Supplementary figure 3.**
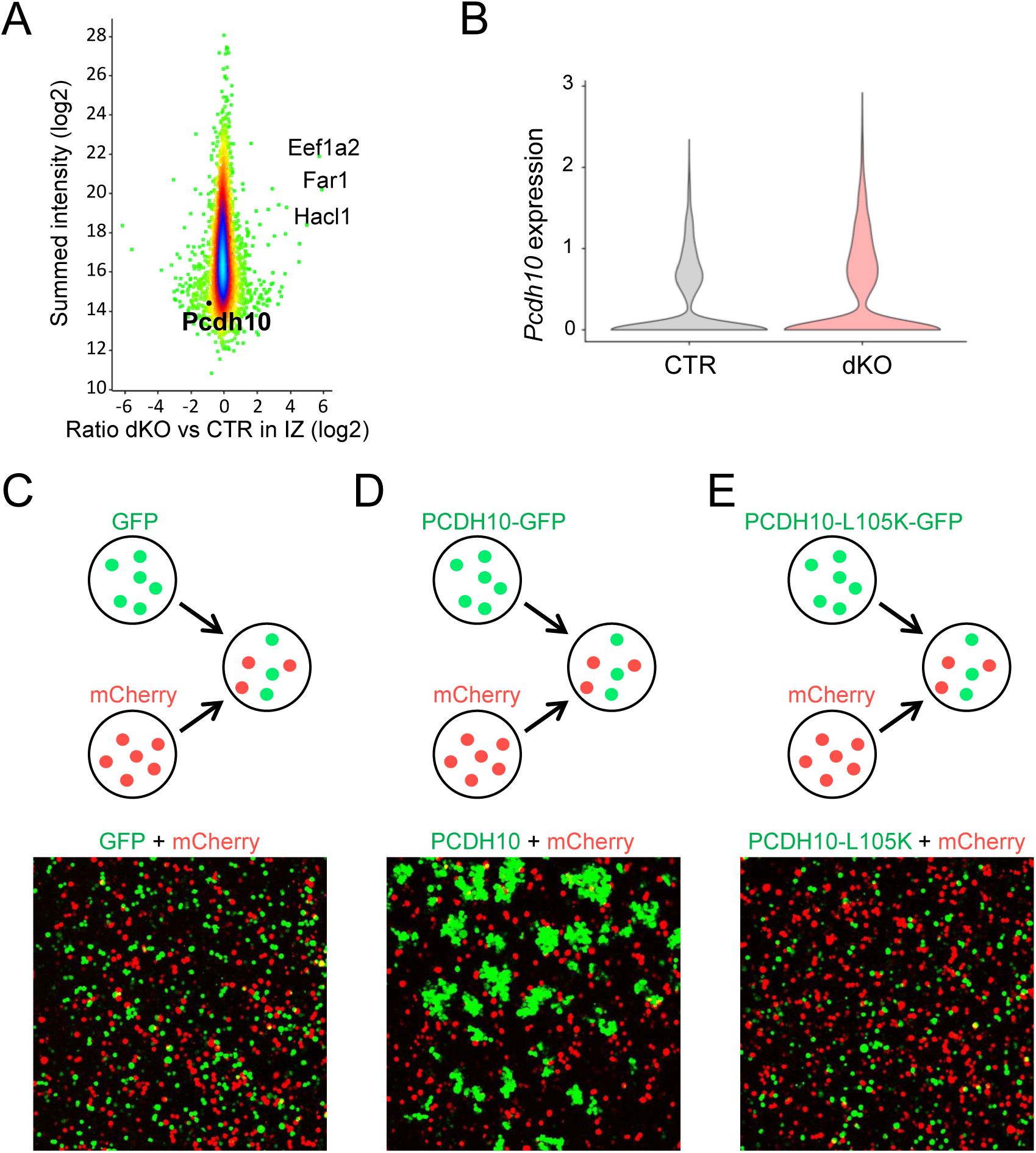
Increased *Pcdh10* expression in *Flrt1/3* dKO and its role in cell aggregation. (A) Volcano plots for IZ show normalized protein expression ratios (dKO vs. CTR, log₂) plotted against summed intensity (log₂). (B) Violin plot showing *Pcdh10* expression in excitatory neurons based on scRNA-seq analysis. (C–E) Cell aggregation assay using K562 cells. Schematics (top) illustrate the mixing of two fluorescently labeled populations. The bottom images show dispersed cell distributions in the GFP control (C) and the PCDH10-L105K mutant condition (E), and aggregation of PCDH10-expressing cells (D).

**Supplementary figure 4.**
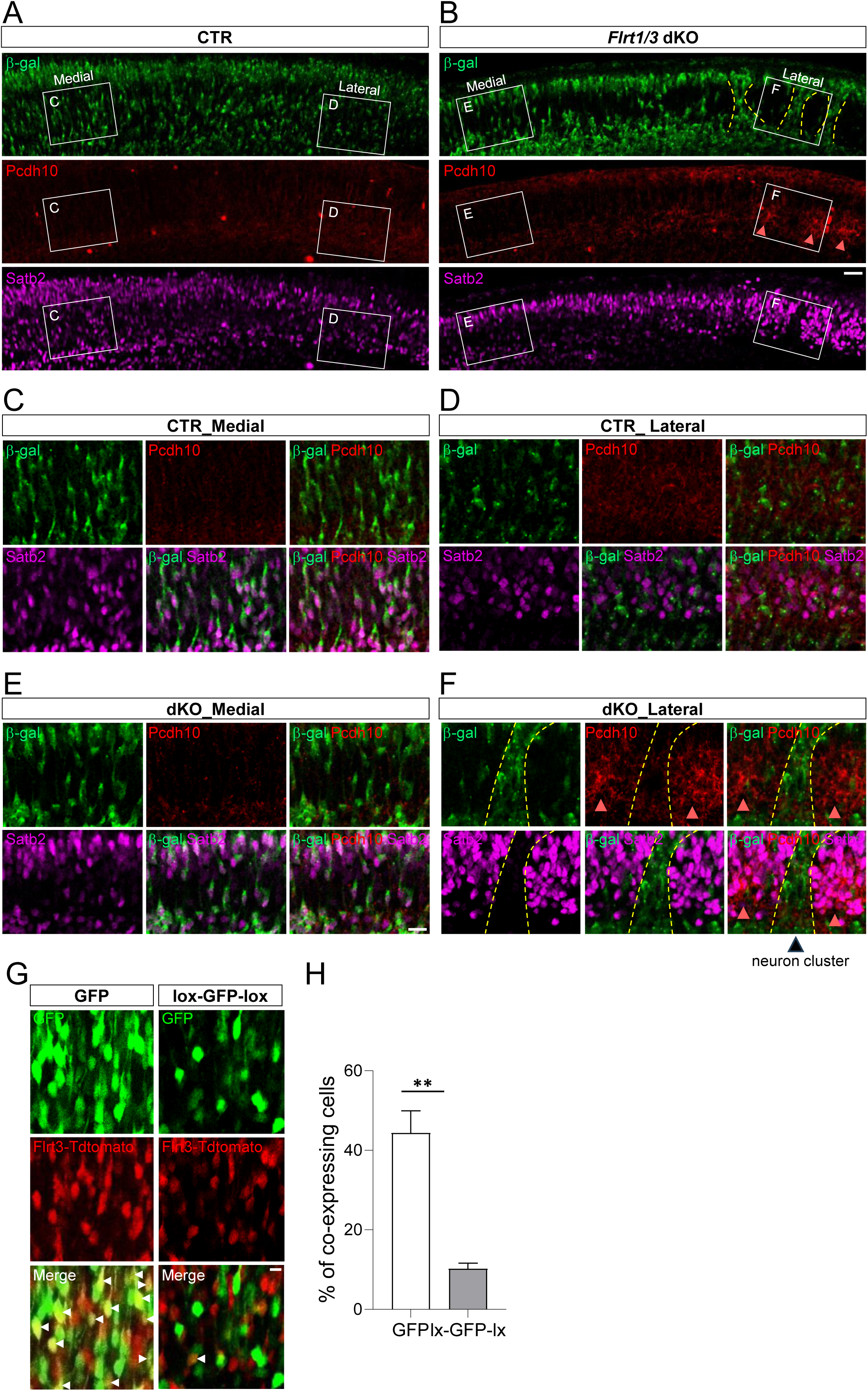
The segregation of *Flrt1/3* mutant clusters and PCDH10+ patches, and validation of genetic strategy for labeling distinct neuronal populations. (A, B) Immunostaining of β-gal (green), Satb2 (magenta), and PCDH10 (red) in the caudal cortex of control (A) and *Flrt1/3* dKO embryos (B). Yellow dashed lines outline clusters of FLRT1/3-deficient neurons, and pink arrowheads mark PCDH10 patches. Scale bar, 50 μm. (C, D, E and F) Higher-magnification images in medial (C and E) and lateral (D and F) portion of control and dKO cortices. Scale bar, 25 μm. (G) Electroporation of GFP constructs with or without loxP sites into *Flrt3-CreERT2/+*; *Ai9* cortices. White arrowheads indicate GFP and tdTomato co-expressing cells. Scale bar, 12 μm. (H) Quantification of co-expressing cells. N = 3 mice per group.

**Supplementary figure 5.**
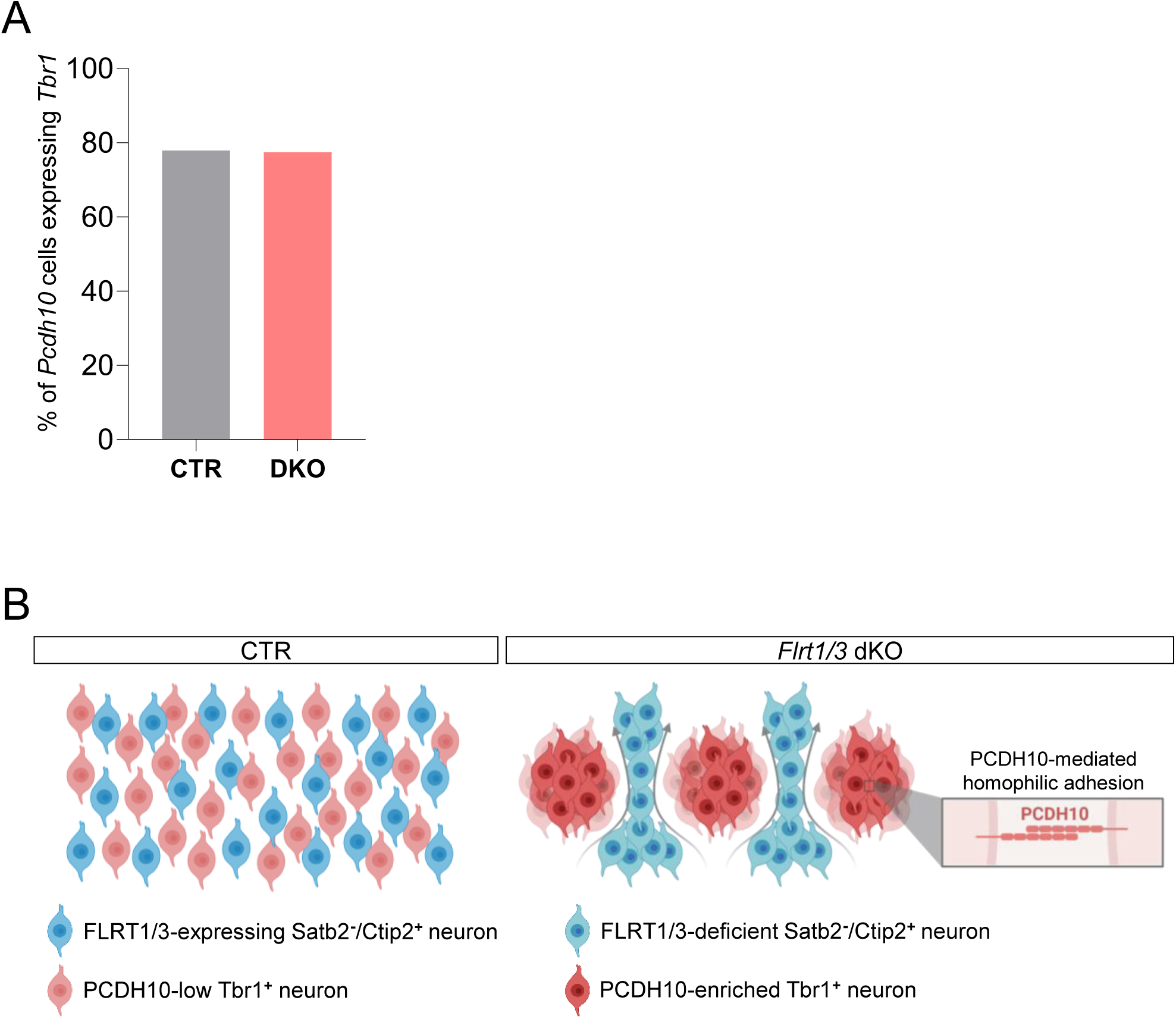
*Pcdh10* expression in Tbr1^+^ neurons and model of PCDH10-mediated spatial segregation of FLRT1/3-deficient neurons. (A) ScRNA-seq analysis shows the percentage of *Pcdh10*-positive excitatory neurons expressing Tbr1. (B) In control cortex, FLRT1/3-expressing Satb2^−^/Ctip2^+^ neurons are intermingled with PCDH10-low Tbr1^+^ cells. In *Flrt1/3* dKO cortex, PCDH10-enriched Tbr1^+^ domains become prominent and, through PCDH10-mediated cell-cell adhesion, act as adhesive compartments that spatially segregate migrating FLRT1/3-deficient Satb2^−^/Ctip2^+^ neurons, leading to their discrete clustering. Created with BioRender.com.

**Supplementary table 1. Differentially expressed genes in excitatory neurons of control versus FLRT1/3-deficient cortices, identified by scRNA-seq analysis.**

**Supplementary table 2. Spatial proteomics of the CP.** Proteins differentially abundant between control and *Flrt1/3* dKO cortices are shown. Significance was assessed using “significance B”. The robust up-regulation of PCDH10 in CP is highlighted in red.

**Supplementary table 3. Spatial proteomics of the IZ.** Proteins differentially abundant between control and *Flrt1/3* dKO cortices are shown. Significance was assessed using “significance B”. PCDH10, highlighted in red, is not significantly changed in the IZ.

## Methods

### Mouse Lines

All animal experiments were approved by the Government of Upper Bavaria (permit numbers 55.2-1-54-2532-57-2015 and ROB-55.2-2532.Vet_02-20-2) and conducted in accordance with the German animal welfare regulations. Mice were housed with 12:12h light/dark cycle in a specific pathogen-free environment with *ad libitum* access to food and water. To generate FLRT1/3-deficient mice, *Flrt3^LacZ/lx^* [53] mice were crossed with *Flrt1^-/-^*[54] and *Nestin-Cre* [55]. To generate *FLRT3-CreERT2*; *Ai9* mice, the *Flrt3-CreERT2* [23] mice were crossed with the *Ai9* [56] mice.

### Single-cell dissociation and RNA sequencing

Brains from E15.5 embryos were sectioned in ice-cold, oxygenated artificial cerebrospinal fluid (ACSF) containing 125 mM NaCl, 26 mM NaHCO_3_, 2.5 mM KCl, 2 mM CaCl_2_, 1 mM MgCl_2_, 1.25 mM NaH_2_PO_4_ (pH 7.4). The medial-to-lateral caudal cortex was micro-dissected and enzymatically dissociated using papain (1 mg/ml, Sigma P4762) in Hibernate E medium supplemented with DNase I (20 µg/ml, Sigma D4513) at 37°C for 15–20 min. Single viable cells were purified by flow cytometry. Dissociated cells from 2 control (*Flrt1^-/-^*;*Flrt3^LacZ/+^*) and 2 dKO (*Flrt1^-/-^*;*Flrt3^LacZ/lox^*;*Nestin-Cre*) samples were prepared for scRNA-seq. A target of 8,000 cells per sample were processed using Single Cell 3’ Reagent Kits (10x Genomics v3.1 chemistry) and libraries were sequenced on an Illumina NextSeq 500.

### Single-cell RNA-seq data processing

Barcode filtering and sequence alignment were performed using the CellRanger pipeline (http://10xgenomics.com). Reads were aligned to the mouse reference genome GRCm38. The gene-by-cell matrices were analyzed using Seurat R package as follows: Cells from control and dKO samples were merged and filtered to retain only higher-quality cells (mitochondrial reads < 7%, genes detected > 200). Next, we normalized the data (normalization.method=’LogNormalize’,scale.factor=10000), detected variable features (selection.method = ’vst’, nfeatures = 3000), scaled the data (vars.to.regress = c(’nCount_RNA’)) and conducted principal component analysis (PCA). We retained 30 principal components for following nearest-neighbor graph construction and UMAP dimension reduction, and unsupervised clustering was conducted at a resolution of 0.8 for entire cell populations and at 0.6 for only excitatory neuron populations.

### 4-Hydroxytamoxifen administration

4-hydroxytamoxifen (Sigma, H-7904) was dissolved in 100% ethanol and diluted in corn oil (Sigma, C8267) to 25 μg/g or 50 μg/g. Pregnant animals with *Flrt3-CreERT2*;*Ai9* embryos were injected intraperitoneally.

### Immunohistochemistry

Brains were fixed in 4% PFA overnight, sectioned and permeabilized in 0.5% Triton X-100, 1% BSA/PBS overnight at 4°C. Sections were incubated with primary antibody: chicken anti-β−galactosidase 1/2000 (Abcam, AB9361), rabbit anti-Tbr1 1/500 (Abcam, AB31940), rabbit anti-Satb2 1/500 (Abcam, AB34735), rat anti-Ctip2 1/1000 (Abcam, AB18465), rat anti-OL-protocadherin 1:1000 (Pcdh10) (Sigma, MABT20), for 48 h at 4°C. The sections were then incubated with secondary antibody Alexa Fluor 488-, 594-and 647-conjugated donkey anti-chicken/rabbit/rat (Molecular Probes 1:500) for 24 h at 4°C. Sections were counterstained with DAPI (Invitrogen, D1306) and imaged using a SP8 laser scanning confocal spectral microscope (Leica Microsystems).

### X-gal staining

Brains were fixed in 1% PFA for 1 h at 4°C and incubated with 30% sucrose in PBS overnight at 4°C. Then brains were embedded in OCT, sectioned at 40 μm, and stained for β-galactosidase activity by incubation at 37°C for 2–3 hr in 1 mg/ml X-gal (Invitrogen) solution containing 5 mM K_4_Fe(CN)_6_ and 5 mM K_3_Fe(CN)_6_.

### Spatial proteomics

Laser capture microdissection (PALM Microbeam, Zeiss) was used to isolate areas of interest from the CP and IZ regions identified by X-gal staining. Collected tissue was lysed in 300 mM Tris/HCl (pH8), 50% 2,2,2-trifluoroethanol for 90 min at 90°C, and sonicated. Proteins were reduced by 5 mM DTT, alkylated by 20 mM CAA, and digested overnight with LysC and trypsin (1:100). Peptides were cleaned by StageTipping and analyzed using LC-MS/MS (Q Exactive HF-X, Thermo Fisher Scientific) coupled to an EASY nLC 1200 ultra-high-pressure system (Thermo Fisher Scientific) via a nano-electrospray ion source. Details of the entire workflow can be found in the recent publication [57]. Data independent acquisition (DIA) raw files were analyzed with the Spectronaut Pulsar X software for direct DIA analysis targeting mouse UniProtKB database (2019). Proteins identified based on a single peptide were filtered out, as well as decoy hits and proteins not passing the default quantification criteria (Q-value cut-off 0.01). For the significance B analysis [58], we calculated an outlier significance score for log protein ratios after intensity binning (at least 300 proteins per bin). Normalised protein ratios from three biological replicates (*Flrt1/3* dKO versus control, log2) were plotted against summed protein intensities (log2). A total of 273 and 314 differentially regulated proteins were identified in CP and IZ, respectively, based on a Benjamini-Hochberg FDR 5%.

### Western blotting

Caudal cortex from control and dKO brains were lysed in buffer containing 50 mM Tris-HCl pH 7.5, 150 mM NaCl, 1% Triton-X 100, 2 mM EDTA, protease inhibitor cocktail (Roche, 04693159001). Protein extracts were resolved on NuPAGE 4-12% gels in SDS-PAGE and transferred onto 0.45 μm PVDF membranes (Merck, IPVH00010). Membranes were blocked in 5 % milk in Tris-buffered saline (TBS; 20 mM Tris-HCl pH 8, 150 mM NaCl) with 0.1% Tween 20 (TBS-T) for 1 h and were incubated with primary antibody: rat anti-OL-protocadherin (pcdh10) antibody (Sigma, MABT20) and mouse anti-β-actin (Sigma, A5316) overnight, at 4°C. After TBS-T washes, HRP-conjugated goat anti-rat (Sigma, A9037) or anti-mouse antibodies (Dako, P0447) were applied for 1 h at room temperature. Blots were acquired using the ChemiDoc MP imager (Bio-Rad).

### Cell aggregation assay

K-562 cells (ATCC CCL-243TM) were transfected with 2.5 μg plasmids using Lipofectamine LTX Reagent (Invitrogen, 11583137) and incubated for 48 hours. Cells were harvested with 0.1 mg/ml DNase I in DMEM containing 5 mM MgCl_2_ and filtered through 40 µm cell strainers to obtain a single cell suspension. Equal numbers of two transfected cell types (1×10⁵ cells each) were co-incubated in Bioinert chambered slides (Ibidi, 80800) for 24 h at 37°C with orbital shaking (80 rpm). Aggregation was imaged by confocal microscopy (Leica SP8).

### *In utero* electroporation

*In utero* electroporation was performed at E12.5 on anesthetized pregnant mice. 1 μl DNA plasmids (1 mg/ml) containing fast green (Sigma, F7252) were injected into the lateral ventricle of mouse embryos. Electric pulses (six pulses, 30 V, 50 ms duration, 1 s interval) were applied using tweezer electrodes. Embryonic brains were harvested at E15.

### Data analysis and statistics

Images were analyzed using software Fiji [59]. Statistical analysis and graphs were generated using GraphPad Prism 9.2.0. Statistical significance was determined using unpaired two-tailed Student’s t-test and was defined as *p < 0.05, **p < 0.01. Data are presented as mean ± SEM.

